# Directionality of two-photon absorption in representative fluorescent proteins

**DOI:** 10.1101/2024.12.31.630429

**Authors:** Josef Lazar, Štěpán Timr, Olga Rybakova, Jiří Brynda, Adam Thuen, Mikhail Drobizhev, Jitka Myšková

## Abstract

Molecules of fluorescent proteins (FPs) exhibit anisotropic optical properties. While the directionality of single-photon (1P) absorption and emission in the most commonly used FPs has already been characterized, directionality of their two-photon (2P) excitation remains poorly understood, even though it offers higher sensitivity to molecular orientation and promises to yield more information on molecular orientational distributions than the 1P processes. By measuring optical properties of FP crystals, we have now determined the two-photon absorptivity tensors (2PATs) of the FPs mTurquoise2, eGFP and mCherry. Our results demonstrate the feasibility of experimental 2PAT determinations, provide new insights into molecular mechanisms of 2P absorption in FPs, open a new avenue to rational development of novel biosensors and allow detailed structural interpretation of results obtained with such biosensors.

## Introduction

Molecules of fluorescent proteins (FPs) exhibit anisotropic optical properties^1^. Optical anisotropy of FPs can be used for detection of molecular interactions, for example, by observations of fluorescence polarization^2^. Anisotropic properties of FP also play an important role in fluorescence resonant energy transfer (FRET), widely used in both in vivo and in vitro biomolecular studies^3^. A thorough knowledge of directional optical properties of FPs allows accurate interpretation of FRET experiments and making detailed insights into molecular structure (precision FRET)^4^. Furthermore, FP optical directionality underpins observations of linear dichroism in living cells, vesicles and other structures, yielding information on cellular division^5^, cell membrane voltage^6,7^, as well as on various molecular processes of cell signaling^8–10^. Uses of FP optical anisotropy are likely to become yet more abundant with recent identification of versatile, widely applicable classes of genetically encoded biosensors (Polaris^11^, FLIPs^12^) that allow facile development of molecular probes for microscopy imaging of a broad range of biomolecular events, even without modifying or overexpressing the protein of interest.

Even though optical anisotropy of FPs has long been known and utilized, its detailed characterization has only been gradually emerging. The directionality of 1P light absorption and emission are characterized by a vector, the transition dipole moment (TDM). The rate of 1P light absorption is proportional to the cos^2^ of the angle between the electric field vector of the incoming photon and the TDM. Similarly, the emission TDM describes the rate of light emission into a particular direction, as well as the polarization of the emitted light. Thus, knowing the orientation of TDMs with respect to the FP atomic framework is crucial for understanding 1P optical directionality of FPs. Although initial studies^1,13^ aiming to elucidate the 1P optical directionality of FPs yielded valuable insights, they suffered from errors in interpretation of molecular structures of FP crystals, as well as in mathematical description. The first determination of TDM orientation in an FP molecule was made on the basis of its relative orientation with respect to the vibration of the carbonyl group in the fluorophore of the green fluorescent protein (GFP)^14,15^. Since then, TDM orientations have been predicted by quantum mechanical calculations for a wide range of FP fluorophores^16,17^. The orientations of TDMs describing both 1P light absorption and emission in mTurquoise2 (mTurq2), eGFP, mCherry and the photoswitchable FP mEos4b have recently been derived by us^18^, by combining optical measurements of FP crystals with detailed information on molecular structures of the crystals. We then complemented this information by computational and experimental investigations^19^ of the dynamics of TDM orientations due to electronic effects, molecular vibrations of the fluorophores and rotational diffusion of FP molecules.

While the directionality of 1P transitions in the most commonly used FPs have by now been characterized both experimentally and computationally, the same cannot be said about two-photon (2P) transitions. Directionality of 2P absorption is described by two-photon absorptivity tensors (2PATs) and can be substantially more complex than directionality of 1P processes^20^. Little experimental information is available on 2PATs of fluorescent molecules. For electronic transitions involving changes in the dipole moment of the fluorophore, 2PATs have been postulated^21^ to be dominated by a single eigenvector with an orientation similar to the orientation of the corresponding TDM. This has generally been confirmed by quantum mechanical (QM) calculations of properties of some fluorescent dyes^22,23^, although with previously unforeseen exceptions^23^. Published QM investigations^24^ of 2P properties of GFP yielded limited agreement with experimental observations and paid no attention to directional aspects. Information on FP TDMs, combined with measurements of fluorescence anisotropy of FP solutions upon 2P excitation with linearly and circularly polarized light has been used to characterize 2PATs in red FPs^25^ (including mCherry) and eGFP. However, more direct and extensive characterization of 2PATs is needed to deepen and solidify our understanding of 2P optical directionality in FPs.

Better understanding of 2P optical directionality in FPs is particularly needed because it promises to allow full exploitation of unique advantages of 2P excitation. As is well known, 2P excitation allows fluorescence imaging deep in living tissues. This is due to the limited dispersion of infrared light used for 2P excitation, as well as due to the inherent optical sectioning ability of 2P excitation. In comparison with 1P excitation, 2P excitation is expected to be markedly more sensitive to molecular orientation (possibly a cos^4^ relationship)^20^. Importantly, linear dichroism observations made by 2P excitation can yield two orientation parameters of the absorbing molecules (such as the mean orientation and the width of the orientational distribution of orientations), providing detailed information not accessible by 1P excitation^26,27^. This information is likely to become indispensable in rational development of new genetically encoded biosensors, as well as in interpreting observations made with such biosensors. The abilities developed for FP-based biosensors will be applicable to small fluorescent labels, including unnatural amino acids, and perhaps even to minuscule molecular and atomic labels for vibrational spectroscopy and microscopy. In order to promote these developments, we have now built upon our experience with TDM determinations through polarization microscopy observations of FP crystals^18^ and carried out extensive characterizations of 2PATs in three widely used FPs: mTurquoise2 (mTurq2), eGFP, and mCherry.

## Results and discussion

### Observations of crystals oriented within the focal plane of the microscope

Our initial measurements of 2P optical properties of FP crystals were performed similarly to our previous determinations^18^ of the directions of FP transition dipole moments (TDMs): on crystals lying flat on the glass coverslip (Fig. 1). However, unlike in our TDM determinations, rather than observing the extent of excitation light transmission, we now measured the intensity of 2P fluorescence as a proxy for 2P absorption. This was because the fraction of illumination absorbed through 2P absorption is minuscule, and thus the intensity of illumination is virtually unaffected by light absorption by the crystal (which hindered using fluorescence intensity for TDM determinations). In order to characterize the transitions into the S1 and S2 excited states, we measured optical properties of each FP at two distinct excitation wavelengths, corresponding to the shorter and longer wavelength absorption bands of the 2P spectra^21^. Crystals of all three FPs exhibited a relationships between laser power and fluorescence intensity that was very close to quadratic, for both investigated wavelengths (Suppl. Fig. 1), confirming the 2P nature of the observed optical phenomena.

**Fig 1:**
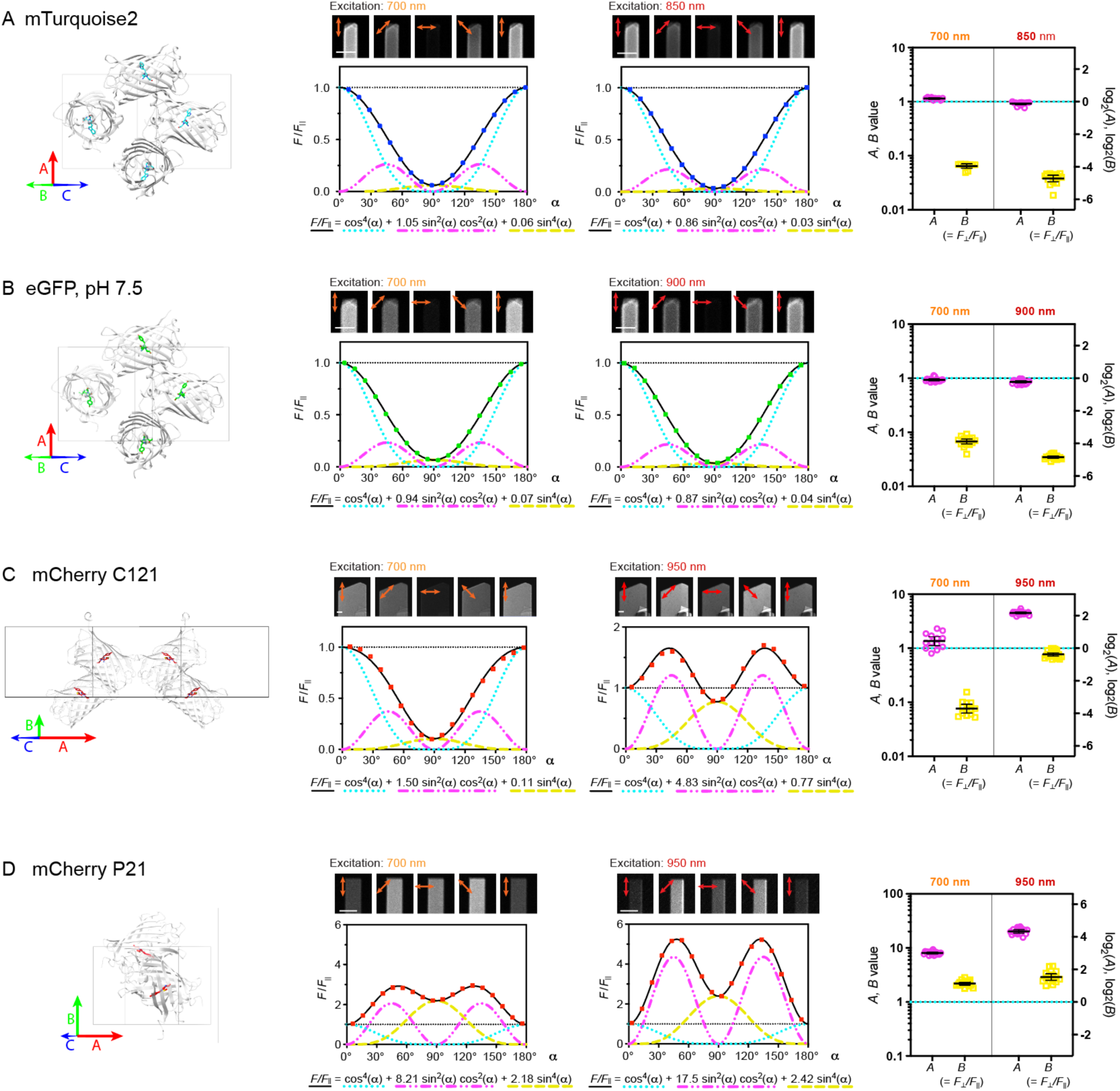
Studied FP crystals and their 2P optical properties observed with crystals oriented within the microscope focal plane. A)-D) Crystals of mTurquoise2 (crystallographic group P212121), eGFP (P212121), mCherry (C121) and mCherry (P21), respectively. Left pane: structure of the crystallographic unit cell, viewed in the direction of the illuminating laser beam, when the crystal is lying flat in the microscope focal plane. The long axis of the crystal is oriented vertically in the shown images. Middle panes: representative images of crystals acquired with different linear polarizations of the excitation light (α = 0°, 45°, 90°, 135°, 180°) and fluorescence intensity profiles (α varied in 10° increments) illustrating data fitting by a cos^4^(α) + *A* sin^2^(α) cos^2^(α) + *B* sin^4^(α) function. Right panes: plots of values *A*, *B* obtained by fitting *F*(α)/*F*(0) profiles. Means ± 2*SEM of Log_2_(*A*), Log_2_(*B*) values are indicated.

Upon illumination with different linear polarizations of the excitation light within the focal plane of the microscope (described by angle α), different types of FP crystals yielded distinct fluorescence intensity profiles (*F*(α)). Crystals of the P212121 symmetry group (mTurq2, eGFP) exhibited markedly more fluorescence (around 20x) when illuminated by light polarized along the long axis of the crystal (α = 0) than by light polarized perpendicular to this axis (α = 90°). The observed fluorescence intensity profiles resembled a cosine squared function and differed only little between the two excitation wavelengths. A similar fluorescence intensity profile was also observed in the mCherry C121 crystals with 700 nm excitation. However, strikingly, with 950 nm excitation, the same mCherry C121 crystals exhibited a very different behavior, with fluorescence intensity minima at excitation polarizations both parallel and perpendicular to the long crystal axis. Two fluorescence intensity minima (at α = 0°, α = 90°) were also observed in mCherry crystals in the P21 crystallographic group, both with 700 nm and 950 nm excitation, although to a different extent with each.

Upon closer inspection, our mathematical model describing 2P absorption by FP crystals revealed that for crystals with a two-fold symmetry axis lying in the focal plane of the microscope, normalized fluorescence intensity profiles can be described by a function in the form *F*(α)/*F*(0) = cos^4^(α) + *A* sin^2^(α) cos^2^(α) + *B* sin^4^(α). Indeed, the observed fluorescence intensity profiles could be well fitted by such a function (Fig 1). Measurements of multiple crystals of each type yielded mean values and 95% confidence intervals of logarithmic values of parameters *A* and *B* (Log_2_(*A*), Log_2_(*B*)) Fig 1, Table 1). The value of parameter *B* describes the ratio of fluorescence intensities observed with excitation polarizations perpendicular and parallel to the long axis of the crystal, respectively (*B* = *F*_⊥_/*F*||).

**Table 1:** Values of coefficients A, B, obtained from measurements on flat-oriented crystals.

| FP, crystal type | Excitation wavelength | $\text{Log}_2(A)$<br>(mean ± 2SEM) | $\text{Log}_2(B)$<br>(mean ± 2SEM) |
| --- | --- | --- | --- |
| mTurq2, P212121 | 700 nm | $0.193 \pm 0.042$ | $-3.95 \pm 0.12$ |
| | 850 nm | $-0.121 \pm 0.056$ | $-4.72 \pm 0.18$ |
| eGFP, P212121 | 700 nm | $-0.098 \pm 0.054$ | $-3.88 \pm 0.14$ |
| | 900 nm | $-0.219 \pm 0.058$ | $-4.84 \pm 0.06$ |
| mCherry, C121 | 700 nm | $0.444 \pm 0.260$ | $-3.71 \pm 0.25$ |
| | 1000 nm | $2.17 \pm 0.05$ | $-0.372 \pm 0.102$ |
| mCherry, P21 | 700 nm | $3.01 \pm 0.05$ | $1.13 \pm 0.09$ |
| | 1000 nm | $4.34 \pm 0.10$ | $1.53 \pm 0.19$ |

### Interpretation of observations of crystals oriented within the focal plane of the microscope using a single eigenvector 2PAT model

In principle, the two experimentally accessible values (parameters *A*, *B*) can be used to determine two parameters of the absorptivity tensors (*S*). Although this is not sufficient for detailed characterization of the absorptivity tensors, it could yield useful information. In order to describe the directionality of 2P excitation by two parameters, we postulated that one of the 2PAT eigenvectors (***l*_1_**) is dominant (the corresponding eigenvalue λ_1_ = 1) while the other two eigenvectors (***l*_2_**, ***l*_3_**) are negligible (λ_2_ = 0, λ_3_ = 0). This allows the directionality of 2P excitation to be characterized by a single vector (***l*_1_**), whose orientation can be described by two parameters: angles τ (orientation of ***l*_1_** within the plane of the fluorophore) and ϕ (tilt of ***l*_1_** with respect to the fluorophore plane). This simplification is supported by the published considerations of relationships between FP 2PATs and TDMs. Furthermore, our observations of 2P fluorescence anisotropy (Suppl. Fig. 2, Suppl. Table 1) suggest that 2PATs of the studied FPs exhibit a single dominant eigenvector whose orientation is close to the emission TDM (β < 20°). To indicate that such 2PATs are defined by a **single** eigenvector we decorate their names with a **single** prime mark (*S*′). Furthermore, to mark the fact that these 2PATs were determined using only data obtained from **horizontally** oriented crystals we use a **horizontal bar** accent 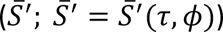.

In order to find tensors 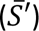 consistent with our observations, we set the eigenvalues (λ_1_, λ_2_ and λ_3_) corresponding to the eigenvectors ***l*_1_**, ***l*_2_**, and ***l*_3_** to 1, 0, and 0, respectively. We then exhaustively sampled the parametric space of angles τ and ϕ, calculating the expected values of Log_2_(*A*) and Log_2_(*B*) for each τ, ϕ combination. The results are summarized in Fig 2, Table 2, and Suppl. Table 2. Briefly, some τ, ϕ pairs matched only the experimentally observed Log_2_(*A*) values, and some only the Log_2_(*B*) values. Finally, some τ, ϕ pairs yielded predictions of both Log_2_(*A*) and Log_2_(*B*) values that fell within the respective experimentally determined confidence intervals.

**Fig. 2:**
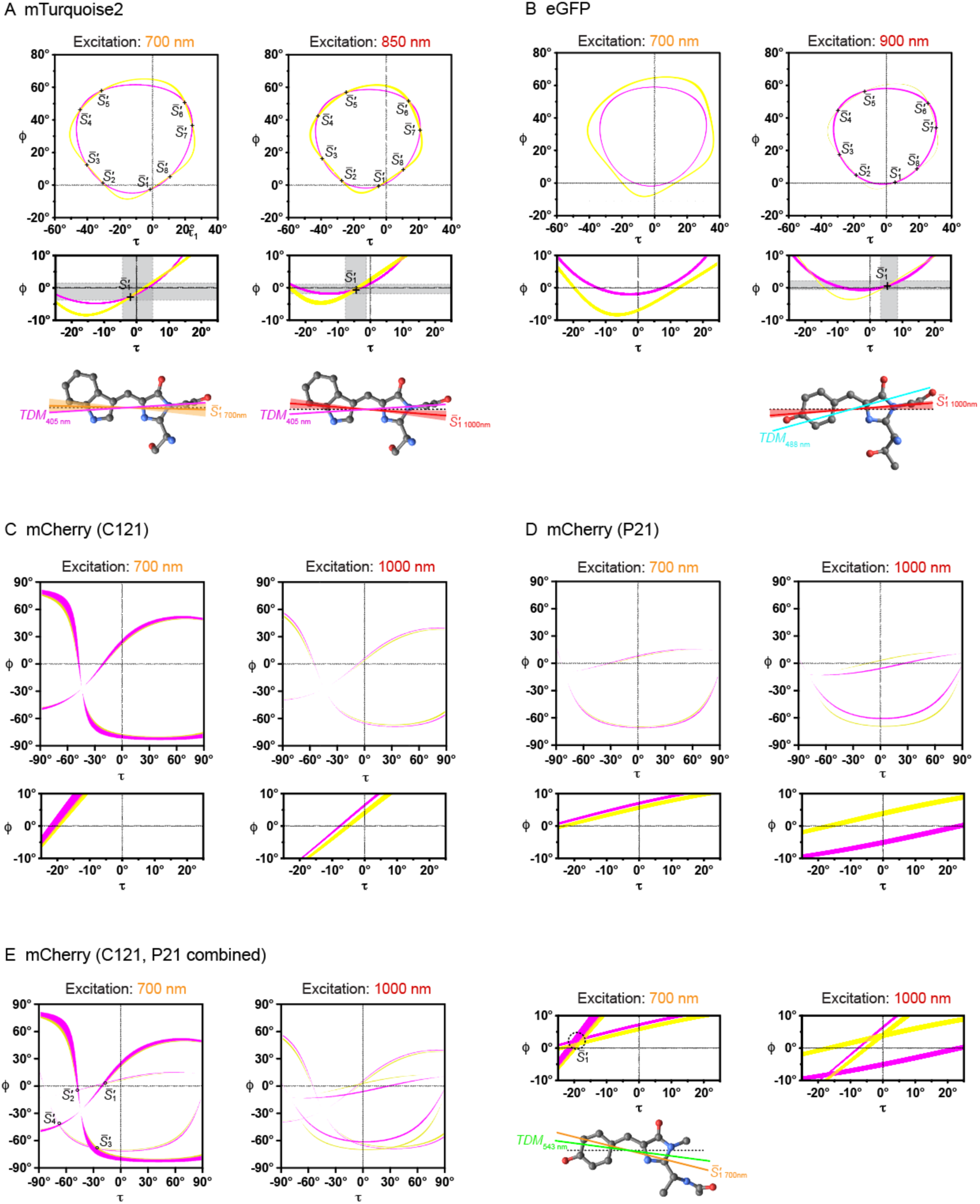
Interpretation of observations of crystals oriented in the focal plane of the microscope, using a single eigenvector 2PAT model. Top: combinations of τ, ϕ values consistent with the observed Log_2_(*A*) and Log_2_(*B*) 95% confidence intervals (in magenta and yellow, respectively). Roots (tensors 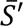) indicated (where applicable). Middle: detailed view of the τ, ϕ space close to the fluorophore plane (small |ϕ|), close to the long axis of the fluorophore (small |τ|). Bottom (where applicable): visualizations (radial plots) of absorption rates for different excitation light polarizations with respect to the fluorophore, for 2PATs [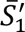]. A) mTurquoise2; B) eGFP; C) mCherry crystals in the C121 group; D) mCherry crystals in the P21 group; E) mCherry, combining data from C121 and P21 crystals.

**Table 2:** Results of 2PAT determinations from FP crystals oriented within the focal plane of the microscope, using a single eigenvector model.

| FP, crystal type | Excitation wavelength | $S$ | $\tau$ [°]<br>mean (95% CI) | $\phi$ [°]<br>mean (95% CI) |
| --- | --- | --- | --- | --- |
| mTurq2, P212121 | 700 nm | $\bar{S}'_1$ | -1.4 (-4.7 – 5*) | -2.7 (-4.2 – 1.1) |
| | 850 nm | $\bar{S}'_1$ | -4.5 (-8.2 – 1.3) | -0.9 (-2.0 – 1.0) |
| eGFP, P212121 | 700 nm | - | - | - |
| | 900 nm | $\bar{S}'_1$ | 4.4 (3.0 – 8.8) | -0.7 (-0.8 – 2.1) |
| mCherry, C121 | 700 nm |  | -90 – 90 | -85 – 85 |
|  | 1000 nm |  | -90 – 90 | -70 – 60 |
| mCherry, P21 | 700 nm |  | -90 – 90 | -75 – 15 |
|  | 1000 nm |  | - | - |
| mCherry, C121, P21 | 700 nm | $(\bar{S}'_1)$ | (-18) | (2) |
|  | 1000 nm |  | - | - |

For crystals of mTurq2 and eGFP (the P212121 space group; Fig. 2A, 2B), the identified τ, ϕ combinations matching experimental observations formed irregular circular shapes in the τ, ϕ plane. In some cases, the τ, ϕ combinations that agreed with the experimentally observed values of Log_2_(*A*) and of Log_2_(*B*) overlapped, yielding absorptivity tensor solutions. This way, 2PATs for mTurq2 (both 700 nm and 850 nm) and eGFP (900 nm only) could be found (Suppl. Table 2). Since this approach yields multiple solutions (typically 8; denoted as 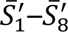), we used the expected similarity between the 2PAT and the known excitation TDM to identify one solution (denoted as 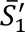) that is the most likely to be the 2PAT (Fig. 2, Table 2). In one instance (eGFP, 700 nm excitation) no τ, ϕ combination was consistent with the experimentally observed values of both Log_2_(*A*) and Log_2_(*B*), suggesting that in this case, approximating the 2PAT by a single eigenvector is not justified.

For crystals of mCherry and 700 nm excitation data, the mathematical model found many τ, ϕ combinations that matched both the observed values of Log_2_(*A*) and Log_2_(*B*) in a particular type of crystal (P21 and C121). This prevented identification of distinct 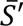 tensors using a single type of crystal, although the data was consistent with the single eigenvector model. However, searching the τ, ϕ space for combinations that would simultaneously agree with Log_2_(*A*) and Log_2_(*B*) observed in both types of mCherry crystals allowed identification four areas nearly matching the experimental observations. We denoted the one closest to the known emission TDM as 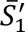 (Fig. 2, Table 2). In contrast with the 700 nm measurements, our observations of mCherry crystals made with 1000 nm excitation could not be satisfactorily interpreted by using the single eigenvector model (Fig. 2).

Thus, our analysis has shown that data from crystals oriented within the focal plane of the microscope is consistent with the single eigenvector model in some cases (mTurq2 both at 700 and 850 nm, eGFP at 900 nm, and mCherry at 700 nm), but not in others (eGFP at 700 nm, mCherry at 1000 nm). When applicable, the model yielded absorptivity tensors 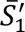 defined by a single vector (***l*_1_**) and angles τ, ϕ, similar to those previously found to describe the corresponding transition dipole moment.

### Observations of crystals rotated along their long axis, and their interpretation using the single eigenvector 2PAT model

To further characterize the directionality of 2P light absorption in the investigated FPs, we performed measurements on crystals rotated along their long axis (Fig. 3). As before, for each observed crystal, the fluorescence intensities measured with different excitation light polarizations (angle α) were fitted by a parametric equation (*F*(α)/*F*(0) = cos^4^(α) + *A* sin^2^(α) cos^2^(α) + *B* sin^4^(α)). Values of the logarithms of the parameters *A*, *B* were then plotted as a function of the crystal roll angle γ (Log_2_(*A*(γ)), Log_2_(*B*(γ))).

**Fig 3.**
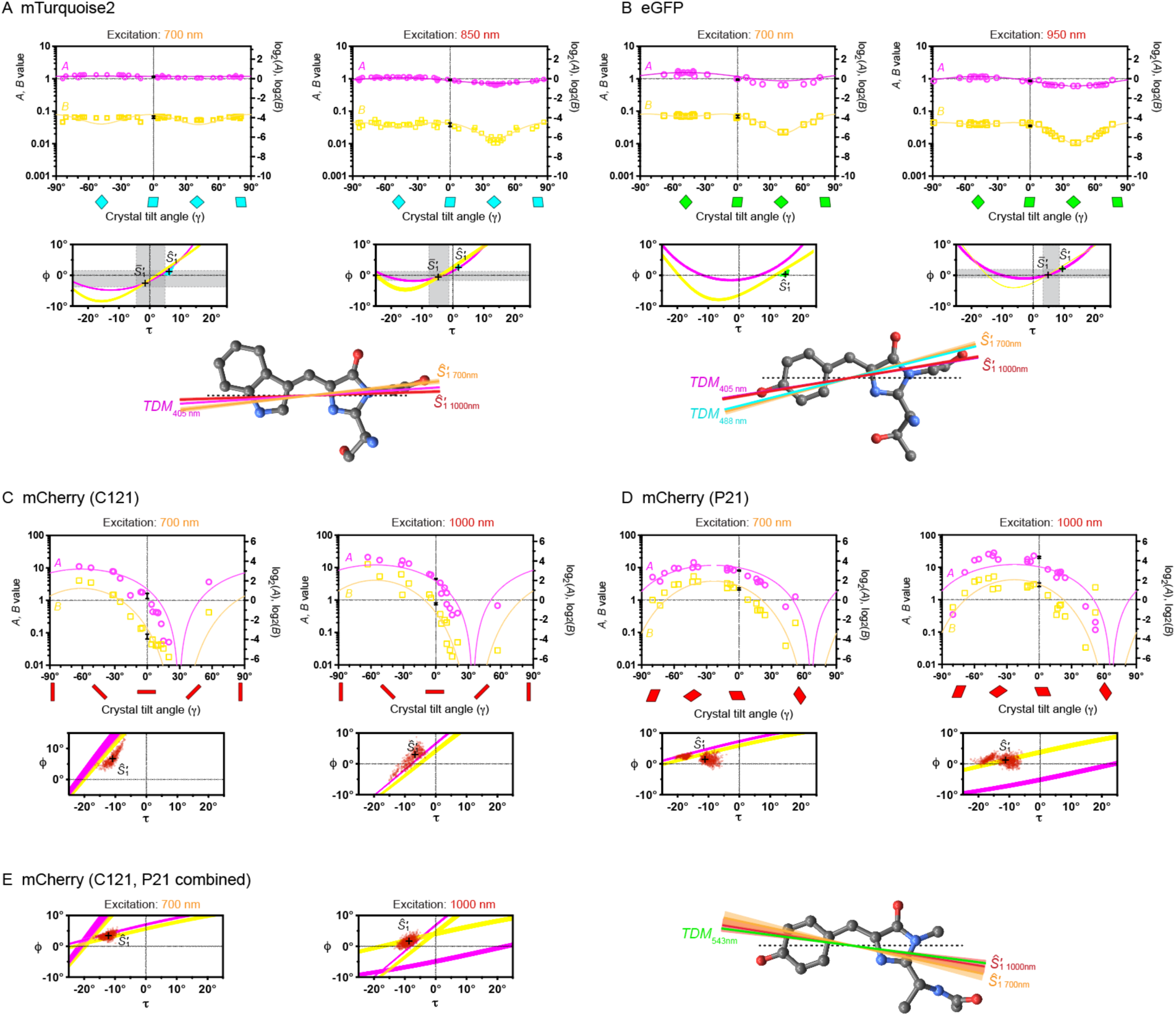
Observations of crystals tilted along their long axis, interpreted using a single eigenvector 2PAT model. A) mTurquoise2; B) eGFP; C) mCherry (C121); D) mCherry (P21); E) mCherry (data from P21 and C121 crystals combined). Larger plots: values of Log_2_(*A*) (magenta circles) and Log_2_(*B*) (yellow squares) observed for different values of the crystal roll angle γ. The 95% confidence intervals of Log_2_(*A*) and Log_2_(*B*) from measurements of crystals oriented flat are marked at γ = 0. Characteristic orientations of the crystal cross-sections are illustrated by colored parallelograms. Best fits of the combined Log_2_(*A*(γ)), Log_2_(*B*(γ)) data, obtained using a single dominant eigenvector 2PAT (*Ŝ′* = *Ŝ′*(τ, ϕ)) model, are indicated by solid lines. Smaller plots: combinations of τ, ϕ agreeing with data obtained using crystals oriented in the focal plane of the microscope (as in Fig 2), along with mean values and 95% confidence intervals of τ, ϕ values obtained from measurements on crystals at different roll angles γ.

Subsequently, the experimentally observed Log_2_(*A*(γ)) and Log_2_(*B*(γ)) relationships were examined in two distinct ways. First, we investigated agreement of observed the Log_2_(*A*(γ)) and Log_2_(*B*(γ)) relationships with predictions made by using 2PATs 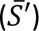, determined as described in the previous section, by observing crystals oriented within the focal plane of the microscope. Next, we tried to identify 2PATs that would agree best with the observed Log_2_(*A*(γ)) and Log_2_(*B*(γ)) data. We denoted such 2PATs 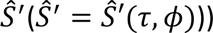, the **hat accent** (instead of a horizontal bar) indicating that **tilted crystals** were used in such determinations, and the **single prime mark** denoting the **single eigenvector** of the mathematical model employed.

Briefly, predictions of Log_2_(*A*(γ)), Log_2_(*B*(γ))) made using 2PATs determined from crystals oriented within the focal plane of the microscope 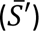 yielded mixed agreement with data obtained using tilted crystals (Suppl. Fig. 3). In mTurquoise2, the Log_2_(*A*(γ)), Log_2_(*B*(γ))) values observed with 700 nm excitation lied clearly outside of the ranges predicted for the 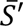 tensors. In contrast, the Log_2_(*A*(γ)), Log_2_(*B*(γ))) values observed with 900 nm excitation in mTurq2 and 950 nm in eGFP lied at the edge of the 95% confidence intervals of predictions made for the respective 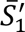 tensors, replicating well with the overall shape of the experimentally observed Log_2_(*A*(γ)), Log_2_(*B*(γ)) relationships. In mCherry, since the crystals oriented in the focal plane of the microscope did not yield a well defined 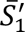 tensor and confidence intervals, a meaningful comparison with data obtained using tilted crystals could not be made. Nonetheless, our results show that the 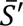 tensors can satisfactorily predict observations of tilted mTurq2 and eGFP crystals excited at their longer wavelength excitation peaks.

Inspired by the apparent ability of the single eigenvector model to reproduce results obtained using tilted FP crystals, we next attempted to find tensors that would yield the best agreement with the observed Log_2_(*A*(γ)), Log_2_(*B*(γ))) relationships (Fig. 3, Table 3, Suppl. Table 3). Simultaneous fitting of the Log_2_(*A*(γ)) and Log_2_(*B*(γ)) data for angles τ, ϕ yielded tensors *Ŝ′*.

**Table 3:** Results of 2PAT determinations from tilted FP crystals, using a single eigenvector model.

| FP, crystal type | Excitation wavelength | $S$ | $\tau$ [°]<br>(mean ± 2SD) | $\phi$ [°]<br>(mean ± 2SD) | $\lambda_1$ | $\lambda_2$<br>(mean ± 2SD) | $\lambda_3$<br>(mean ± 2SD) |
| --- | --- | --- | --- | --- | --- | --- | --- |
| mTurq2,<br>P212121 | 700 nm | $\hat{S}'_1$ | $6.3 \pm 0.7$ | $0.9 \pm 0.6$ | 1 | 0 | 0 |
| | 850 nm | $\hat{S}'_1$ | $1.8 \pm 0.3$ | $2.6 \pm 0.3$ | 1 | 0 | 0 |
| eGFP,<br>P212121 | 700 nm | $\hat{S}'_1$ | $15.1 \pm 0.9$ | $0.4 \pm 0.6$ | 1 | 0 | 0 |
| | 900 nm | $\hat{S}'_1$ | $9.5 \pm 0.2$ | $2.1 \pm 0.2$ | 1 | 0 | 0 |
| mCherry,<br>C121 | 700 nm | $\hat{S}'_1$ | $-11.1 \pm 3.3$ | $6.7 \pm 3.9$ | 1 | 0 | 0 |
| | 1000 nm | $\hat{S}'_1$ | $-7.7 \pm 5.5$ | $2.0 \pm 5.5$ | 1 | 0 | 0 |
| mCherry,<br>P21 | 700 nm | $\hat{S}'_1$ | $-12.7 \pm 7.9$ | $1.8 \pm 2.0$ | 1 | 0 | 0 |
| | 1000 nm | $\hat{S}'_1$ | $-12.8 \pm 7.3$ | $1.5 \pm 2.0$ | 1 | 0 | 0 |
| mCherry,<br>C121, P21 | 700 nm | $\hat{S}'_1$ | $-12.2 \pm 2.6$ | $3.5 \pm 1.5$ | 1 | 0 | 0 |
| | 1000 nm | $\hat{S}'_1$ | $-9.0 \pm 2.9$ | $1.6 \pm 2.1$ | 1 | 0 | 0 |

In mTurq2 and eGFP, for each excitation wavelength, four such tensors 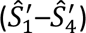 related by crystallographic symmetries of the P2121212 space group were identified (Suppl. Table 3). The identified *Ŝ′* tensors provided identical predictions of Log_2_(*A*(γ)) and Log_2_(*B*(γ)) relationships, in excellent agreement with our observations for the whole range of crystal tilt angles γ. Predictions of Log_2_(*A*(γ)) and Log_2_(*B*(γ)) for γ = 0 lied only slightly outside of the 95% confidence intervals of values observed in crystals oriented within the focal plane of the microscope (Fig. 1). Among the identified symmetry related *Ŝ′* tensors, the one whose dominant eigenvector was closest (within 10°) to the previously determined emission TDM vector was denoted as 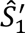 (defined by angles τ, ϕ) and considered to be the 2PAT (Fig. 3, Table 3). Robustness of our determinations was investigated by bootstrapping, yielding standard deviations for τ, ϕ smaller than 0.5°.

In mCherry crystals (both P21 and C121), fitting of Log_2_(*A*(γ)) and Log_2_(*B*(γ)) data for angles τ, ϕ yielded pairs of tensors 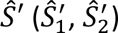 related by crystallographic symmetry (Suppl. Table 3). Of each pair, the tensor whose dominant eigenvector was closer to the previously determined emission TDM was denoted as 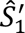 and considered to be the 2PAT (Fig. 3, Table 3). Notably, the Log_2_(*A*(γ)) and Log_2_(*B*(γ)) data for mCherry is more noisy than in mTurq2 and eGFP, leading to larger standard deviations (up to 4°) in our determinations of τ and ϕ. This is partly due to the values of Log_2_(*A*(γ)) and Log_2_(*B*(γ)) being very small in some mCherry crystal orientations, and therefore hard to accurately determine. Furthermore, in mCherry crystals, the values of Log_2_(*A*(γ)) and Log_2_(*B*(γ)) appeared to be quite sensitive even to small deviations of the long crystal axis from horizontal direction. Finally, the shapes of the mCherry C121 crystal cross-sections do not allow reliable determinations of the sign of the tilt angle γ. Because of these factors, 2PAT determinations from a single type of mCherry crystals are markedly less precise than for mTurq2 or eGFP. Combined fitting of data from both types of crystals improves precision of 2PAT determination from measurements of tilted crystals, yielding a dominant eigenvector orientation close to the previously determined emission TDM. However, the agreement of the resulting predictions of Log_2_(*A*(γ)) and Log_2_(*B*(γ)) for γ = 0 with our observations of crystals oriented in the focal plane of the microscope is only limited. Thus, while both shorter and longer wavelength 2P absorption bands of mTurq2 and eGFP can be accurately described by the single eigenvector model, this is less true for mCherry.

## Conclusions

Here we present the results of our characterizations of directionality of 2P optical properties in representative fluorescent proteins. By utilizing a combination of X-ray crystallography and optical measurements by 2P polarization microscopy, we were able to show that mTurq2 and eGFP can be well described by a 2P with a single dominant eigenvector, which we term characteristic eigenvector. The orientations of the characteristic eigenvectors are close (within 5°) of the orientations of the previously determined TDMs of 1P excitation. The eigenvectors describing the S_0_ → S_1_ and S_0_ → S_2_ transitions differs by only a few degrees.

Unlike in mTurq2 and eGFP, crystals of mCherry exhibited striking differences in optical properties between the two 2P absorption bands (700 and 1000 nm). The 2P absorption in mCherry crystals could be described less well by the single eigenvector model than mTurq2 and eGFP. However, the deviations did not appear dramatic for either excitation wavelength. Despite the marked differences in optical properties of mCherry crystals between the two wavelengths used, the orientations of the characteristic eigenvectors differ only by 5°. This is illustrative of the high sensitivity of 2P absorption to excitation light polarization (cos^4^ relationship for the single eigenvector model; Fig. 4).

**Fig. 4.**
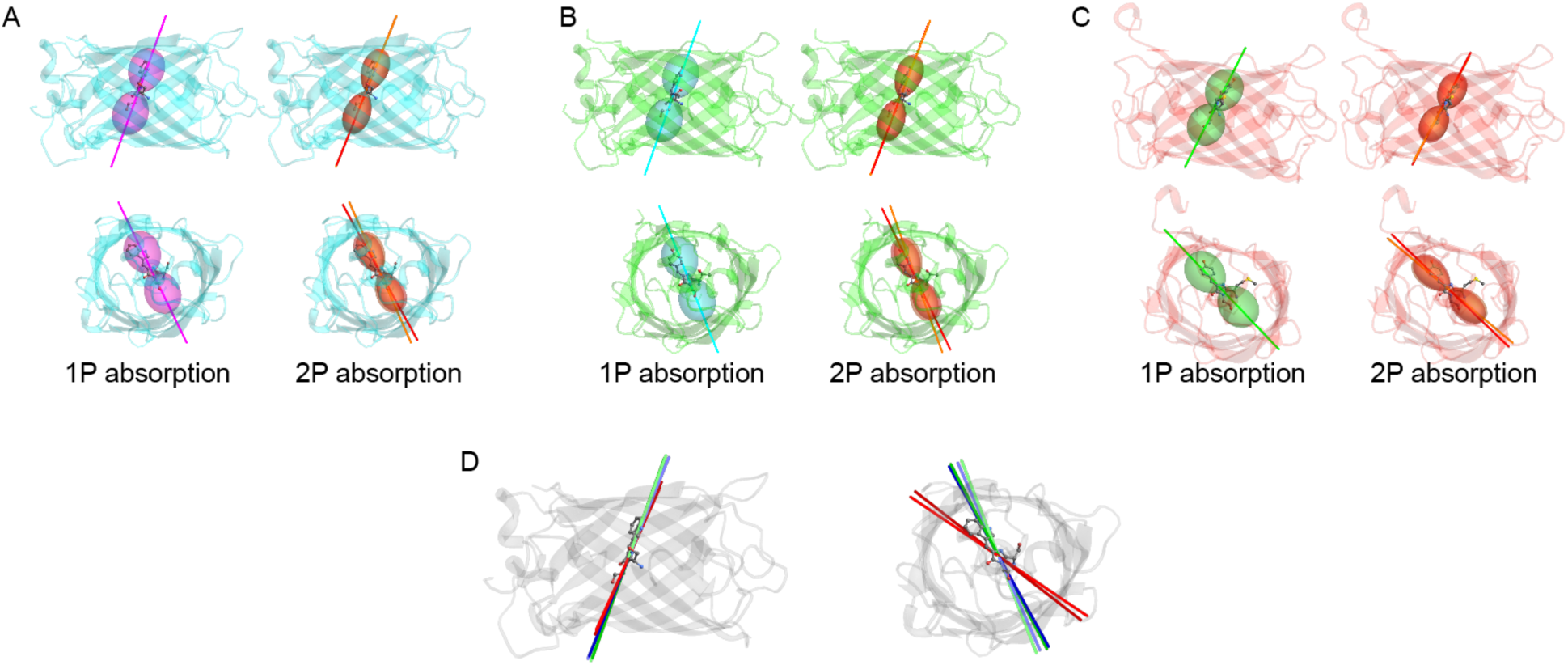
Comparison of directionality of 1P and 2P absorption in mTurq2 (A), eGFP (B) and mCherry (C). The colored lines indicate the published directions of 1P TDMs^18^ and the dominant eigenvectors of 2PATs [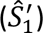] for the shorter- and longer-wavelength absorption band of each FP (in orange and red, respectively). The colored lobes are radial diagrams, indicating the relative rates of light absorption for different polarizations of the excitation light. The larger width of 1P diagrams is indicative of a less pronounced polarization selectivity (cos^2^ relationship) than in 2P absorption (cos^4^ relationship). D) Comparison of orientations of the dominant eigenvectors of the [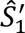] 2PATs with respect to the FP β-barrel, after alignment to the structure of mTurq2. Colors indicate the FP: mTurq2 (blue), eGFP (green), mCherry (red). Lighter shade corresponds to the shorter wavelength absorption band. Lateral and axial views are shown throughout.

Our characterizations of 2PATs in mTurq2, eGFP and mCherry fill an important gap in our knowledge of directionality of light absorption in FPs. Our experimental confirmation of the single eigenvector model validates our assumptions in previous tentative determinations of FP orientations with respect to the cell membrane^26^. Knowledge of the characteristic eigenvector orientations will make these determinations more precise. Furthermore, measurements of linear dichroism of cellular compartments bearing FP-labeled proteins at distinct excitation wavelengths might allow determinations not only of the orientation of the characteristic eigenvector, but of the FP β-barrel. This, in turn, will yield valuable structural insights into function of genetically encoded biosensors, as well as various signaling proteins, directly in living cells, in real time. The expertise and tools gained will be transferable to smaller fluorescent labels (including unnatural fluorescent amino acids) and to other optically active moieties. Thus, our characterizations of FP 2PATs potentially open a new path towards detailed characterization of protein structures in living cells. Apart from unlocking this possibility, our results also provide a verification of our combined crystallography/polarization microscopy approach, enhancing the credibility of our previous determinations of TDMs. The approach, including the mathematical tools we developed, will allow detailed characterization of other optical properties of biomolecules.

## Acknowledgments

Funding was provided by the Czech Science Foundation grant 23-05983S (to J.M.) and NIH/NINDS grant 2U24NS109107 (M.D.). We thank P. Khoroshyy for technical assistance and for maintenance of microscopy equipment. We learned from discussions with P. Jungwirth. We appreciate the contribution of numerous participants of the annual International Summer School in Molecular Biophysics in Nové Hrady, particularly F. Batysta, K. Poncová, Y. Mayr and E. Kot. We are grateful to K. Tošnerová for technical assistance.

## Author contributions

J. Lazar constructed optical equipment, performed initial crystal measurements, developed image processing software tools, performed mathematical modeling and data analyses, prepared manuscript figures and contributed to manuscript writing. Š. Timr carried out optical measurements, as well as performed and supervised mathematical modeling. O. Rybakova carried out microscopy measurements, crystallographic analyses and mathematical modeling. J. Brynda performed crystal structure determinations and assisted in optical measurements. A. Thuen carried out measurements of fluorescence anisotropy. M. Drobizhev supervised and carried out measurements of fluorescence anisotropy, analyzed data, and contributed to writing the manuscript. J. Myšková supervised the project, purified and crystallized fluorescent proteins, performed microscopy observations, image processing and analyses, contributed to writing, and finalized the manuscript.

## Conflicts of interest

J. Lazar is a founder and owner of Innovative Bioimaging, s.r.o. and Innovative Bioimaging, L.L.C, companies manufacturing and selling equipment for polarization-resolved fluorescence microscopy. Both companies may indirectly benefit from the described work, although neither company financially contributed to it.

## Experimental

### Microscopy

Crystals of FPs were prepared as described before^18^, using the hanging drop method. Prior to microscopy, crystals were placed in a drop of mother liquor and sandwiched between two glass coverslips sealed by an adhesive spacer (SecureSeal, Grace Biolabs). Microscopy observations were carried out on a modified laser scanning inverted microscope Olympus FV1200. A tunable femtosecond laser MaiTai HP 1040 (Newport/SpectraPhysics, USA) attenuated to output power suitable for crystal observations (around 10 mW) was used for excitation. The linear polarization of the laser beam was purified by a Glan laser-polarizing beamsplitter (GL10B, Thorlabs, Germany), prior to entering the microscope scanning unit.

During measurements of polarization-dependent optical properties of crystals, polarization of the excitation light was rotated by 720° in 10° increments by a super-achromatic half-wave plate (SAHWP05M-700, Thorlabs), mounted in a motorized rotating mount (PRM1/MZ8, Thorlabs) in a compartment directly below the objective turret. The crystals were illuminated through a low NA objective lens (UPlanFLN 10x, NA 0.3, Olympus). Fluorescence intensity was measured in the trans direction, by an external photomultiplier mounted above the sample. Upon entering the detector unit, laser light was removed by optical filters. For assessment of crystal shape and orientation, image z-stacks of 1-μm spacing were acquired using a 40x objective lens (UApoN 340W, 40x, NA 1.15, Olympus). For each type of crystal and excitation wavelength we measured at least 10 crystals positioned flat on the surface of the coverslip and 10 crystals tilted along their long axis.

### Microscopy data processing

Microscopy images were processed in Fiji, using standard tools and in-house developed macros, in a fashion similar to that used by us previously. Attention was paid to potential sources of systematic errors and other factors that might negatively affect the accuracy of our determinations, particularly as the observed crystals often exhibited large polarization-dependent changes in fluorescence intensity, which needed to be quantitated accurately.

Briefly, series of images acquired with different excitation light polarizations (orientations of the half-wave plate) were aligned using the StackReg plugin. Background was subtracted using an in-house developed ImageJ macro that guided the user to select 3 distinct background areas, whose pixel coordinates (*x*, *y*) and intensities (*z*) were then used to define a plane, which was subtracted from the whole image. Finally, an image area corresponding to a flat part of the crystal free of discernible defects was manually selected, and the intensity of this area was measured for each image within an image series. These intensities were then used for further analysis.

Our mathematical model (see below) revealed that, for crystals with an axis of two-fold rotational symmetry lying within the focal plane of the microscope, the relationship between 2P fluorescence intensity (proportional to the 2P absorption rate) and the polarization of the excitation light within the focal plane (defined by the angle α) can be described as *F*(α) = *k* (cos^4^(α) + *A* sin^2^(α) cos^2^(α) + *B* sin^4^(α)). In order to eliminate effects of differences in crystal sizes, as well as to equalize the statistical weight of the data corresponding to fluorescence maxima and minima, we normalized the intensities *F*(α) across each image series, and calculated logratios of the normalized fluorescence intensities. To overcome issues with precisely aligning the observed crystals with the direction of excitation polarization, we did not normalize by using values of *F*(0°), *F*(90°), *F*_max_ or *F*_min_, but rather by *F*(α + 90°). Thus, measurements of each crystal yielded a series of values of log_2_(*F*(α)/*F*(α + 90°)). Fitting by log_2_[(cos^4^(α) + *A* sin^2^(α) cos^2^(α) + *B* sin^4^(α))/(cos^4^(α + 90°) + *A* sin^2^(α + 90°) cos^2^(α) + *B* sin^4^(α + 90°))] then yielded the values of parameters *A* and *B* associated, respectively, with the sin^2^(α) cos^2^(α) and cos^4^(α) parts of the *F*(α) description. Statistical analysis of Log_2_(*A*) and Log_2_(*B*) obtained from multiple crystals allowed us to obtain 95% confidence intervals for both Log_2_(*A*) and Log_2_(*B*). Using logarithmic values (Log_2_(*A*), Log_2_(*B*)) allowed us to transform a ratiometric meaning of *A*, *B* into a logratio, suitable for Gaussian statistics. Furthermore, it allowed fair comparisons of confidence intervals across a wide range of values of *A* and *B*.

### Mathematical model

Mathematical modeling and absorptivity tensor determinations were performed in Mathematica (Wolfram Research). A coordinate system of the microscope (*x* and *y* oriented horizontally and vertically in the image plane; *z* corresponding to the optical axis of the system) was used throughout. The polarization of the excitation light was described by an electric field vector ***e***(α):

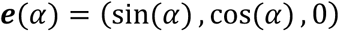

The FP crystals were modeled to be oriented with the long axis of the crystal (a two-fold crystallographic symmetry axis) coinciding with the *y* axis and a flat face of the crystal lying in the *xy* plane. This arrangement allowed using the atomic (pdb) coordinates obtained through X-ray crystallography, either without any transformation, or after a suitable rotation along the *x* or *y* axis (by an angle γ_1_ or γ_2_, respectively), by applying the corresponding rotation matrix. As in our previous studies, each investigated fluorophore was geometrically approximated by a plane identified by least-square fitting of the coordinates of heavy atoms participating in the conjugated bond system. The long axis of the fluorophore was defined as a line connecting the centers of the aromatic rings.

The absorptivity tensor (*S*) was generically defined by a set of mutually orthogonal, unit-sized eigenvectors (***l*_1_**, ***l*_2_**, ***l*_3_**) and the respective eigenvalues (λ_1_, λ_2_, λ_3_). We described the orientation of this eigensystem by angles τ (rotation along the fluorophore plane normal) and ϕ (rotation along the short axis of the fluorophore), so that for τ = 0 and ϕ = 0, ***l*_1_** and ***l*_2_** would correspond to the long and short axes of the fluorophore, respectively, and ***l*_3_** would be the normal of the fluorophore plane. Thus, the absorptivity tensor *S* was described by 5 parameters (τ, ϕ, λ_1_, λ _2_, λ_3_):

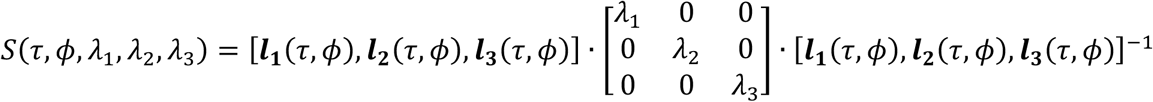

Since we normalized the experimentally observed crystal fluorescence intensities, the eigenvalues of the absorptivity tensor *S* could also be normalized, by setting the value of λ_1_ to 1:

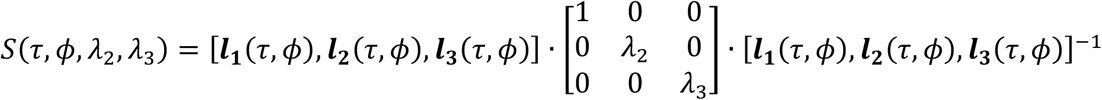

This normalization reduced the number of parameters to 4. Assumptions about the orientation of the eigensystem (such as setting ϕ = 0) and/or dominance of a single eigenvector (that is, setting λ_2_ = λ_3_ = 0) allowed simplifying the model to match the number of parameters yielded by experimental data.

To account for the multiple orientations of the FP molecules within an FP crystal, a set of symmetry-related absorptivity tensors (*S*_*sym1*_, *S*_*sym2*_,…, *S*_*symn*_) was generated for each molecular orientation within a given crystal type, by applying the relevant symmetry operation matrices ([*sym_1_*], [*sym_2_*],…, [*sym_n_*]):

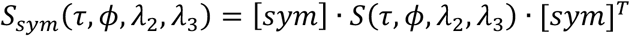

For the P212121 crystallographic group (mTurquoise2, eGFP), we used the following symmetry matrices:

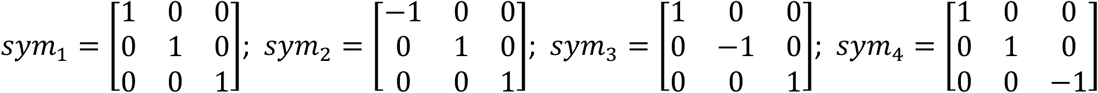

For the P21 and C121 crystallographic groups (mCherry), both with two-fold symmetry along the crystallographic axis B, we used the following symmetry matrices:

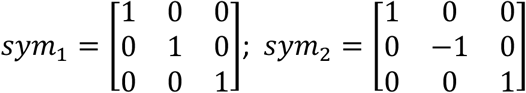

The resulting set of symmetry-related tensors was oriented within the microscope coordinate system in a way simulating the crystal positioned on a microscope slide by applying the rotation matrix [*rot*(γ)]. This allowed us to predict the fluorescence intensity of a crystal for different crystal orientations, absorptivity tensors, and excitation light polarizations (electric field vector ***e***(α)):

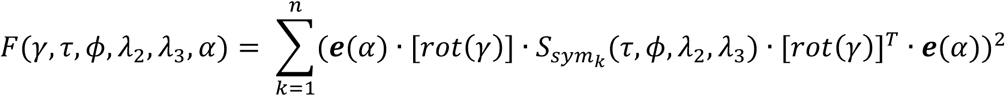

For a given combination of tensor parameters (τ, ϕ, λ_2_, λ_3_) and crystal orientation (γ), the mathematical model yielded values of fluorescence intensity as a function of the excitation polarization angle (α), in the form *F*(α) = *k* (sin^4^(α) + *A* sin^2^(α) cos^2^(α) + *B* cos^4^(α)). Comparing the values of *A* and *B* predicted for a particular combination of γ, τ, ϕ, λ_2_ and λ_3_ with the experimentally determined values of *A* and *B* then allowed us to find the absorptivity tensors that matched our experimental data.

### Absorptivity tensor determinations

In order to determine the absorptivity tensors using the single eigenvector model, we set the values of λ_2_, λ_3_ to be equal to zero. We identified the τ, ϕ combinations matching the observed values of Log_2_(*A*), Log_2_(*B*) (tensors *S*^′^) by sampling values of the two angles over the interval <-90°, 90°> in 0.1° increments. We used the 95 % confidence intervals of Log_2_(*A*), Log_2_(*B*) to identify the confidence intervals of τ, ϕ.

Tensors *Ŝ′* were identified by fitting the measured Log_2_(*A*(γ)) and Log_2_(*B*(γ)) data by the mathematical model described above. By varying τ, ϕ, we minimized the sum of root mean square deviations (RMSDs) from both experimental sets of values. For mCherry, the sum of RMSDs from the observed Log_2_(*A*(γ)) and Log_2_(*B*(γ)) data for both types of crystals (C121 and P21) was used to find angles τ, ϕ that would best match the combined data. As minimization starting points we used the τ, ϕ values obtained in our *S*^′^determinations. Standard deviations of τ, ϕ were estimated by 100 rounds of bootstrapping.

### Other optical measurements

Measurements of 2P fluorescence anisotropy were carried out as before^25^, on purified FPs used for crystallizations. Coefficients *r*_0_, *r*_1_ were calculated using a previously published procedure.^28^ Dependence of fluorescence intensity on excitation laser power was measured using FP crystals using a microscopy arrangement similar to that used in other FP crystal observations. Laser power was controlled by an attenuator consisting of a manually rotatable achromatic half-wave plate and a Glan-laser polarizer. Laser power was measured using a photodiode sensor (Thorlabs, S121C) and a power meter (Thorlabs, PM100D).

---

**Suppl. Fig. 1:**
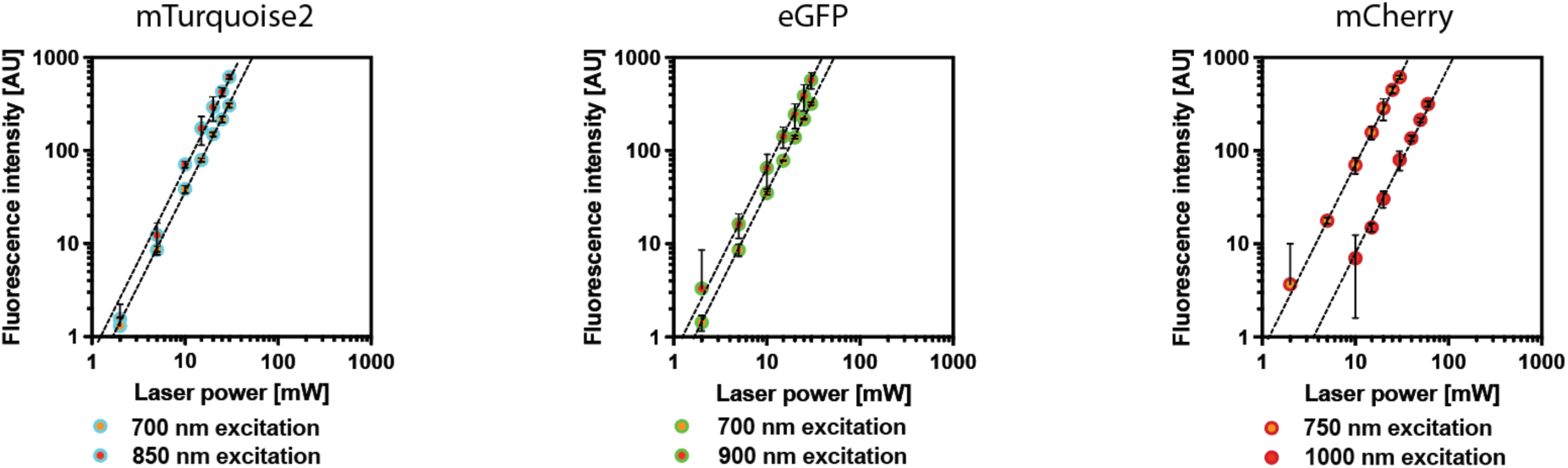
Relationships between excitation laser power and observed fluorescence intensities. Error bars indicate 95% confidence intervals. Dotted lines indicate a quadratic relationship (slope = 2.0).

**Suppl. Fig. 2:**
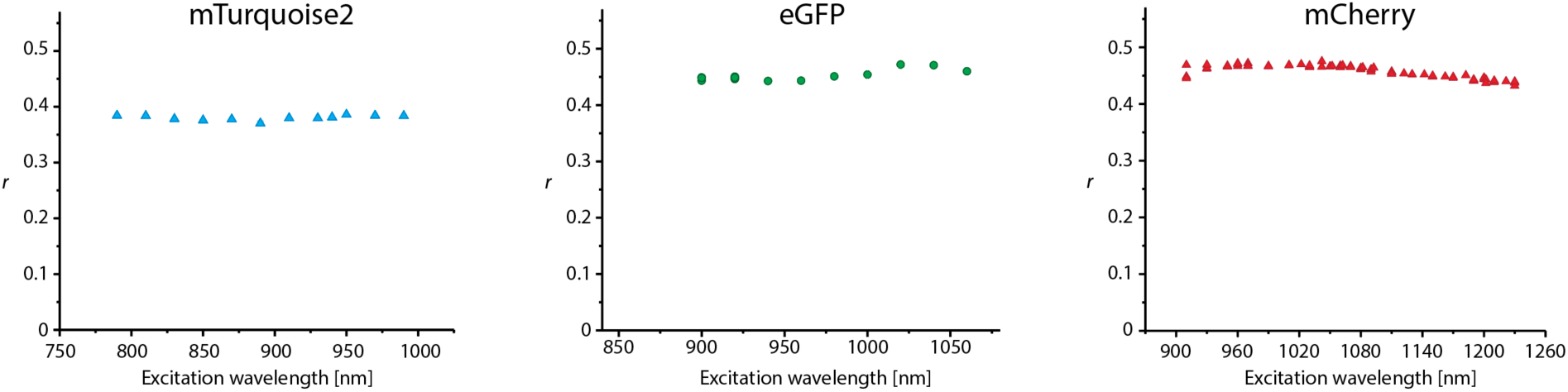
Two-photon excited fluorescence anisotropy measured in FP solutions

**Suppl. Fig. 3:**
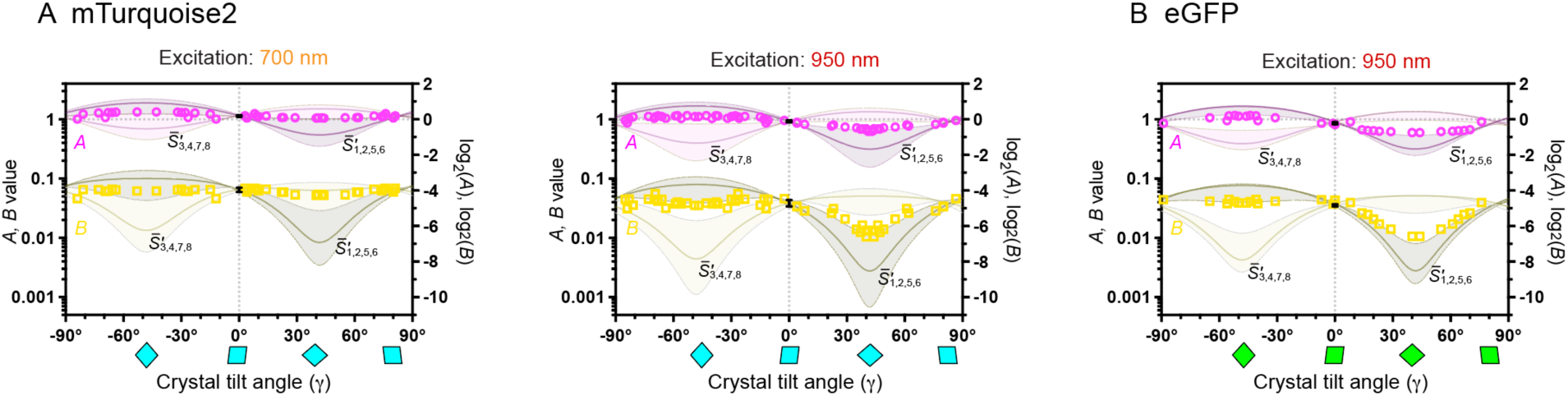
Predictions of Log_2_(*A*(γ)), Log_2_(*B*(γ))) for 2PATs determined by using crystals oriented in the focal plane of the microscope (tensors *S*^′^). A) mTurquoise2; B) eGFP.

**Suppl. Table 1:** Interpretation of 2P fluorescence anisotropy observations.

| FP | Excited state lifetime | Fluorescence anisotropy ($r$) | Initial fluorescence anisotropy ($r_0$) | $\beta$ |
| --- | --- | --- | --- | --- |
| mTurquoise2 | 4.0 ns | 0.38 | 0.48 | 20° |
| eGFP | 2.6 ns | 0.45 | 0.52 | 14° |
| mCherry | 1.4 ns | 0.44 | 0.48 | 18° |

**Suppl. Table 2:** Results of 2PAT determinations from flat oriented FP crystals, using a single eigenvector model.

| FP, crystal type | Excitation wavelength | $S$ | $\tau$ [°]<br>mean (95% CI) | $\phi$ [°]<br>mean (95% CI) |
| --- | --- | --- | --- | --- |
| mTurq2,<br>P212121 | 700 nm | $\bar{S}'_1$ | -1.4 (-4.7 – 5*) | -2.7 (-4.2 – 1.1) |
| | | $\bar{S}'_2$ | -30.6(-35* – -27.9) | 1.4 (-0.9 – 5.8) |
| | | $\bar{S}'_3$ | -40.3 (-42.3 – -35*) | 12.3 (5.7 – 15.4) |
| | | $\bar{S}'_4$ | -44.4 (-46.1 – -41.2) | 45.8 (42.3 – 50*) |
| | | $\bar{S}'_5$ | -31.0 (-41.6 – -25.5) | 57.9 (50* – 60.1) |
| | | $\bar{S}'_6$ | 19.6 (17.0 – 22.9) | 50.9 (44* – 54.2) |
| | | $\bar{S}'_7$ | 24.3 (22.8 – 24.7) | 36.7 (32.9 – 44*) |
| | | $\bar{S}'_8$ | 10.7 (5* – 13.5) | 5.4 (1.0 – 7.9) |
| | 850 nm | $\bar{S}'_1$ | -4.5 (-8.2 – 1.3) | -0.9 (-2.0 – 1.0) |
| | | $\bar{S}'_2$ | -27.1 (-32.0 – -23.9) | 2.3 (0.0 – 6.8) |
| | | $\bar{S}'_3$ | -39.3 (-41.4 – -35.2) | 16.2 (10.5 - 20) |
| | | $\bar{S}'_4$ | -42.0 (-43.6 – -37.5) | 42.2 (38.2 – 48.6) |
| | | $\bar{S}'_5$ | -24.6 (-33.1 – 17.9) | 56.8 (52.4 – 58.7) |
| | | $\bar{S}'_6$ | 13.9 (10.1 – 18.1) | 51.4 (45.2 – 54.7) |
| | | $\bar{S}'_7$ | 20.9 (19.8 – 21.5) | 33.8 (29.6 – 40.6) |
| | | $\bar{S}'_8$ | 10.8 (5.3 – 13.7) | 9.4 (4.8 – 12.4) |
| eGFP,<br>P212121 | 700 nm | - | - | - |
| | 900 nm | $\bar{S}'_1$ | 4.4 (3.0 – 8.8) | -0.7 (-0.8 – 2.1) |
| | | $\bar{S}'_2$ | -18.2 (-21.2 – -16.9) | 3.4 (2.9 – 7.5) |
| | | $\bar{S}'_3$ | -29.9 (-30.0 – -26.2) | 17.8 (13.8 – 19.2) |
| | | $\bar{S}'_4$ | -31.3 (-31.3 – -26.5) | 43.7 (42.3 – 48.0) |
| | | $\bar{S}'_5$ | -12.0 (-18.3 – -9.5) | 57.5 (54.0 – 57.7) |
| | | $\bar{S}'_6$ | 25.7 (24.2 – 27.4) | 50.4 (45.6 – 51.3) |
| | | $\bar{S}'_7$ | 31.4 (29.8 – 30.9) | 32.5 (31.3 – 37.8) |
| | | $\bar{S}'_8$ | 20.2 (15.3 – 20.7) | 8.8 (6.1 – 10.0) |
| mCherry,<br>C121 | 700 nm |  | -90 – 90 | -85 – 85 |
|  | 1000 nm |  | -90 – 90 | -70 – 60 |
| mCherry,<br>P21 | 700 nm |  | -90 – 90 | -75 – 15 |
|  | 1000 nm |  | - | - |
| mCherry,<br>C121, P21 | 700 nm | $(\bar{S}'_1)$ | (-18) | (3) |
| | | $(\bar{S}'_2)$ | (-49) | (-2) |
| | | $(\bar{S}'_3)$ | (-32) | (-63) |
| | | $(\bar{S}'_4)$ | (-65) | (-42) |
|  | 1000 nm |  | - | - |

\* An approximate value, as the confidence interval overlaps with a confidence interval of another solution

**Suppl. Table 3:**
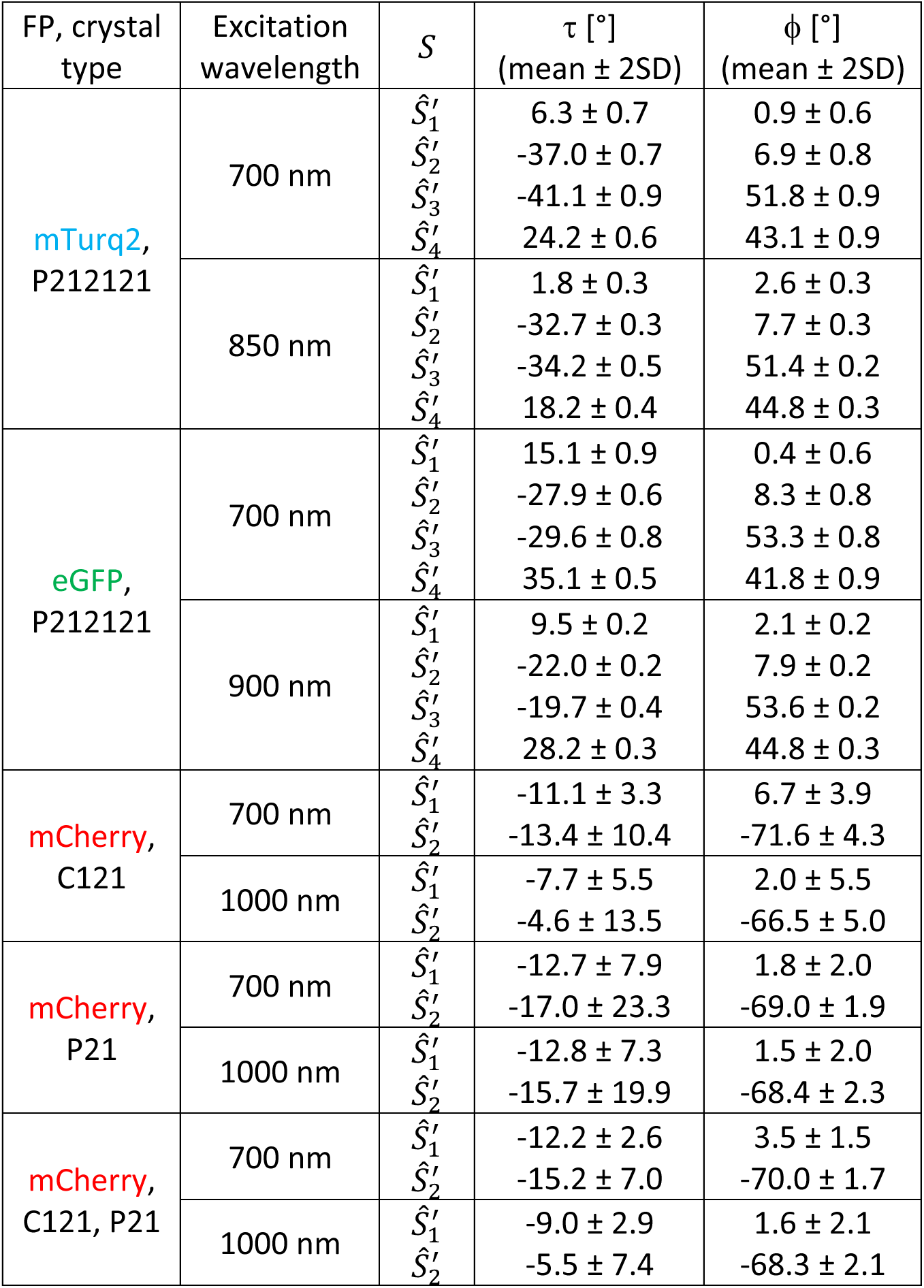
Results of 2PAT determinations from tilted FP crystals, using a single eigenvector model.

## Notes

### Competing Interest Statement

J. Lazar is the CEO of Innovative Bioimaging s.r.o. and Innovative Bioimaging L.L.C. P. Miclea is a part-time employee of Innovative Bioimaging s.r.o. Both companies might indirectly benefit from the results presented herein, although neither company contributed financially or otherwise to the presented research.

